# PCLIPtools: A Robust Framework for Identifying RNA-Protein Interaction Sites from PAR-CLIP experiments

**DOI:** 10.1101/2025.11.17.688892

**Authors:** Ahsan H. Polash, Markus Hafner

## Abstract

PAR-CLIP is a widely used method for identifying binding sites of RNA-binding proteins (RBPs) transcriptome-wide. A characteristic T-to-C transition in the sequenced cDNA pinpoints the site of RBP-RNA crosslinking and is induced by the use of a photoreactive uridine analogue, 4-thiouridine (4SU). As with other systems-wide methods, PAR-CLIP, too, is prone to false discoveries as the T-to-C signal might result from systematic noise, pre-existing SNPs, and PCR errors. Therefore, rigorous statistical methods are required for analyzing PAR-CLIP data. The few existing tools to analyze PAR-CLIP data lack updates and sufficient documentation, and often fail to process current higher-depth sequencing data. Here we report PCLIPtools, a lightweight, customizable suite for analyzing PAR-CLIP data. PCLIPtools considers the read depth, T-to-C transitions, and the other mutations to statistically estimate high-confidence interaction sites. Benchmarking shows that PCLIPtools identifies more functionally significant targets than the current standard tool, PARalyzer, without losing high-confidence sites and outperforming it in runtime. Exploratory analyses show PCLIPtools’ specific targets are enriched for read depth and T-to-C conversion, supporting their validity. With simplicity, robustness, and speed, PCLIPtools improves the precision of PAR-CLIP data analysis while being accessible to experimental RNA biologists. PCLIPtools can be found on github (https://github.com/paulahsan/pcliptools).

**Graphical Abstract:** 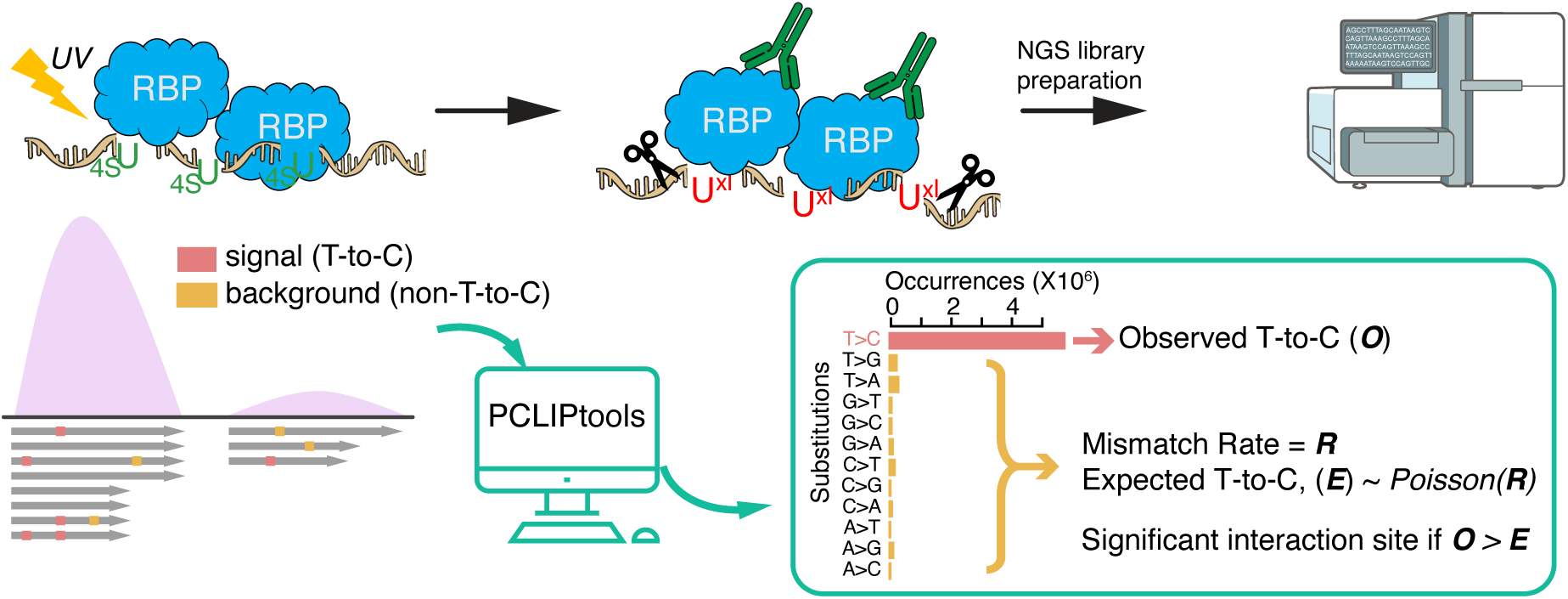

## Introduction

Gene expression in eukaryotic systems is tightly regulated and involves multiple checkpoints and processing steps from RNA biogenesis to translation. On chromatin, precursor mRNAs (pre-mRNA) undergo initial processing, such as capping, splicing, and cleavage and polyadenylation, leading to the formation of mature mRNA (1,2). Additionally, base modifications are installed on all four nucleobases both co- and post-transcriptionally. Mature mRNAs need to be exported from the nucleus to the cytoplasm and correctly localized to specific subcellular compartments for translation (1,2). Ultimately, the mRNAs are degraded as part of the natural turnover process. These various steps are coordinated and controlled by RNA-binding proteins (RBPs), which are proteins that interact with mRNA and non-coding RNAs. To gain fundamental insight into gene regulatory processes, it is crucial to understand the molecular determinants of interactions between RNA and RBPs. This task is complicated by the large complement of RBPs in a eukaryotic cell, their generally high expression levels, and their dynamic localization. Furthermore, while RBPs bind to RNA in a sequence- and structure-dependent manner, the typical RNA binding domain (RBD) of an RBP recognizes short sequences (3,4), resulting in hundreds or even thousands of different RNAs that can be bound and potentially regulated by a single RBP. As a result, systems-wide approaches are necessary to experimentally identify RBP targets and their binding sites to characterize posttranscriptional regulatory processes.

To date, several high-throughput techniques have been developed to study RNA-binding protein (RBP)-RNA interactions by purifying the ribonucleoprotein complexes assembled by RBPs and bound RNAs. These techniques include RIP (RNA immunoprecipitation) (5) and, now more commonly, CLIP (Crosslinking and Immunoprecipitation) (6,7), along with its main variants: eCLIP (8), iCLIP (9), iCLIP2 (10), irCLIP (11), and PAR-CLIP (3,12). All CLIP methods depend on the irreversible covalent crosslinking of RNA with interacting RBPs through irradiation with UV light (13). This covalent linkage of RNA and protein enables stringent purification of the RNA-protein complexes, typically via immunoprecipitation, followed by limited fragmentation of the RNA to isolate the sites occupied by the RBP. High-throughput sequencing of the RNA fragments then permits the identification of interaction sites of the RBPs on a transcriptome-wide scale using computational analysis that typically involves some form of peak calling.

The different experimental approaches offer unique advantages and limitations. Here, we focus on PAR-CLIP (3,12), which relies on metabolic labeling of nascent RNA in living cells with 4-thiouridine (4SU). 4SU-labeled RNA efficiently photo-crosslinks to interacting RBPs when exposed to UVA or UVB light (λ > 312 nm). Photo-crosslinking causes a change in the base-pairing properties of 4SU, leading it to preferentially pair with guanosines during reverse transcription, resulting in a characteristic T-to-C mutation in the sequenced cDNA library. Quantifying these T-to-C transitions simplifies data analysis, as these mutations indicate *bona fide* interaction sites at nucleotide resolution. Furthermore, their enrichment may help assess the strength of the RNA-RBP interaction. Nevertheless, given the vast size of the genome, most next-generation sequencing (NGS) datasets are susceptible to capturing background noise (14,15). The empirical probability of any random genomic region aligning NGS reads with enrichment of T-to-C mismatches is quite low and therefore, PAR-CLIP-derived RBP interaction sites exhibit reduced noise and lower false discovery rates, making it a reliable technique for studying RBP-RNA interactions.

Since PAR-CLIP differs from the rest of the CLIPs due to the T-to-C mismatch signature, the rationale for its peak calling method differs from those used in other CLIP techniques. To accurately predict statistically significant RNA-RBP interaction sites, robust statistical measures must be utilized that account for both read alignment and T-to-C enrichment. Despite the availability of multiple tools for calling PAR-CLIP peaks, only a few (e.g., PARalyzer (16), wavClustR (17,18), BMix (19)), and PARA-suite (20) were specifically developed for PAR-CLIP. Notably, only one of them, PARalyzer, is widely used for scientific discovery based on the number of publications claiming to use it (**Fig. 1A**). CLIP Tool Kit (CTK) (21) and omniCLIP (22) have also been used for analyzing PAR-CLIP data (**Fig. 1A**), although both of them are general peak callers for several forms of CLIP data (13,21). A significant limitation is that many of these PAR-CLIP-specific tools are developed by computational groups that do not conduct PAR-CLIP experiments themselves and therefore benchmark with previously published, low-depth data. Additionally, some tools were developed nearly a decade ago and have not received any updates recently, lacking support from the developers and often missing proper documentation to run them effectively. Due to recent advances in small RNA library preparation and sequencing technologies, current NGS libraries are deeper than ever, posing an extra burden for these tools, which often require long processing times (19) and sometimes fail to generate proper logs for troubleshooting, creating a gap between the newest PAR-CLIP data and the analysis methods available.

**Figure 1.**
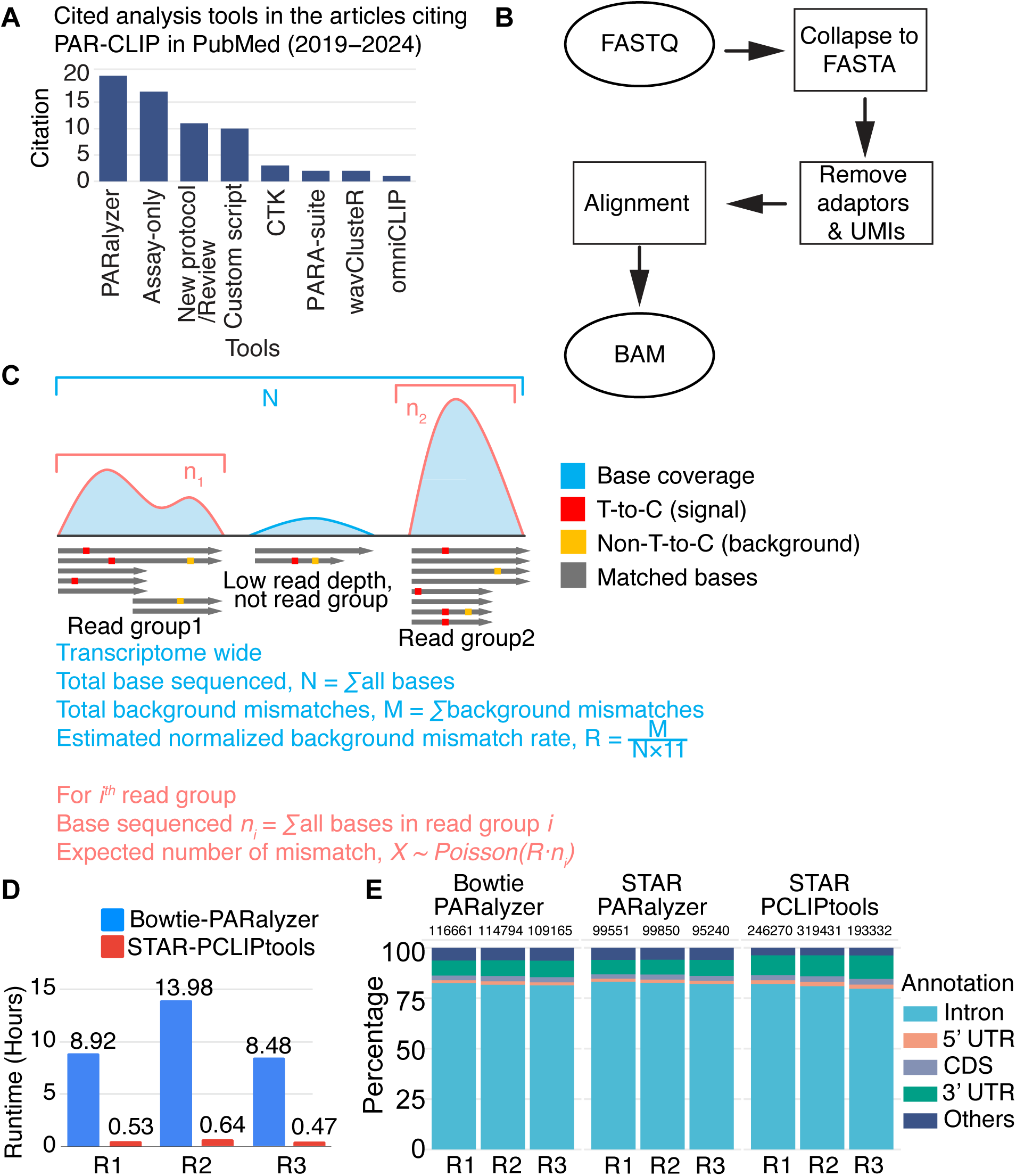
(**A**) PARalyzer is the most widely adopted tool for PAR-CLIP data analysis. (**B**) PCLIPtools employes a simple yet robust statistical approach to align the raw data and (**C**) call peaks. (**D**) PCLIPtools outperfroms widely used tool PARalyzer in runtime (**E**) by retaining intron as major binding site for MATR3. R1, R2, R3 depicts three replicates; UTR, untranslated regions.

Each peak caller uses a unique and robust statistical method to identify RBP-RNA interaction sites. For example, PARalyzer employs a Gaussian kernel density-based classification method to differentiate T-to-C mismatch events from T-to-T matches. A significant concern in NGS data is the presence of mismatches introduced during library preparation via reverse transcription and PCR, base-calling errors, and single-nucleotide polymorphisms (SNPs) (17). In practice, most PAR-CLIP peak callers ignore these sources of mismatches, increasing the risk of false discoveries. Algorithmically, wavClusteR (18) and BMix (19) provide greater robustness compared to other tools, as they employ mixture models that incorporate additional mismatch sources, including non-T-to-C mismatches, SNPs, and more, to detect RNA-RBP interaction sites. However, the respective publications for each tool reported re-analyses of older PAR-CLIP data, which do not fully demonstrate the potential of these tools with current, deeper datasets. Most importantly, none of these tools offer complete pipelines and lack sufficient documentation to resolve running errors. For instance, wavClusteR occasionally fails with certain BAM files generated by STAR (23) aligners and provides inadequate logs for further troubleshooting (the author closed the project in October 2023 https://github.com/FedericoComoglio/wavClusteR/issues/10#issuecomment-1756982601). The installation of PARalyzer, CTK, BMix, and OmniCLIP is also cumbersome due to their reliance on multiple external libraries, which often break and make them difficult to set up and use on high-performance computational clusters (HPCs). Consequently, PAR-CLIP data analysis necessitates an updated tool that employs robust statistical methods, addresses all types of mismatch sources, and is easy to install and operate, with minimal external library requirements. Additionally, the tool should include basic scripts for visualizing PAR-CLIP analysis results.

Most of the existing tools that process PAR-CLIP data use separate scripts for execution and defining operational parameters, compelling users to manually edit configuration files before each run, which reduces the program’s flexibility and adaptability. These limitations represent a hurdle to the adoption of PAR-CLIP (or other CLIP methods) by laboratories that do not have specialized computational expertise and are studying the molecular biology and genetics of RBPs. Therefore, we aimed to develop a tool that is both easy to install and execute, without dependencies on external programs with specific version requirements. Additionally, our tool should support command-line parameters and allow users to run batch processes with a single script. We present a new peak calling tool for PAR-CLIP, called PCLIPtools, which features a comprehensive pipeline for analyzing PAR-CLIP data. The peak caller completes the analysis with higher confidence and reduced processing time compared to other available tools. It models various types of background mismatch events by fitting to a Poisson distribution to calculate significant T-to-C signal enrichments. This facilitates the discovery of a more accurate transcriptome-wide RNA-RBP interaction map. Additionally, the suite includes utility scripts for performing quality control of the CLIP experiments and comparing results with biological replicates and PAR-CLIP data from other RBPs. The peak calling program and pipeline are written in Bash, resulting in minimal dependencies on external libraries. Consequently, it is easy to install, and aside from core computational expertise, users with limited shell experience will be able to effectively use and modify the tool to suit their needs.

## Materials and Methods

### Data source

We downloaded the raw FASTQ files for all PAR-CLIP data from the Short Read Archive (24) by using the ‘fasterq-dump’ program of the SRA-toolkit package (https://github.com/ncbi/sra-tools). This program facilitates multicore/parallel download of the data, thus reducing time. The following SRA IDs were used for MATR3 PAR-CLIP data: SRR28479449, SRR28479450, and SRR28479451 (25). For DHX36 (26) and Nuclear HNRNPK (27), PAR-CLIP SRA ids were SRR6191036 and SRR23199821, respectively. These raw FASTQ files were investigated for quality check by running FastQC v0.12.1 (https://www.bioinformatics.babraham.ac.uk/projects/fastqc/). After ensuring that the reads are suitable for further analyses, they were aligned to the human genome, and peak calling was performed by using PCLIPtools (https://github.com/paulahsan/pcliptools) and PARalyzer (https://ohlerlab.mdc-berlin.de/software/PARalyzer_85/).

### Alignment and Peak calling by PCLIPtools

PCLIPtools has two main modules: ‘pcliptools-align’ and ‘pcliptools-cluster’. As the names suggest, the first module aligns the FASTQ to the provided genome and generates the BAM file. The second module uses the BAM file to call the peaks and predict the high-confidence interactions. In the following subsection, we briefly discuss the procedures.

### Processing, mapping of short-read data

Depending on the compression of the input FASTQ files, ‘pcliptools-align’ first decompresses and extracts the data. Reads are then collapsed into FASTA format by removing duplicates and retaining only those with a minimum read length of 13 nucleotides. After removing the duplicates, cutadapt (version 4.5) (28) is used to remove the 3’ adapters, 5’ adapters, and unique molecular identifiers (UMIs) (**Fig. 1B**). An additional step is performed to remove reads less than 13 nucleotides. The resulting FASTA file is saved in compressed FASTA format. Notably, adapter sequences, length of UMI, and minimum length threshold parameters can be supplied dynamically via command-line parameters.

In the subsequent steps, the compressed FASTA files are fed into the STAR aligner to map the reads to the hg38 reference genome, or a user-supplied genome build. Since we removed the adapters prior to the alignment, we perform an ‘end-to-end’ alignment mode for STAR. Some of the crucial parameters for STAR were ‘–outFilterMultimapNmax 10’ that limits the maximum number of multi-mappers allowed for a read to 10. The maximum number of mismatches is set to 2 to match the parameters recommended and used in PARalyzer, and therefore, ‘ --outFilterMismatchNmax’ was set to 2.

To complete the alignment process, STAR breaks down the reference genome into shorter indices and splits the reads into smaller subsequences called seeds. These seeds serve as anchors to initiate the alignment procedure. Therefore, length and placement of the seeds are crucial for sensitive alignment, particularly for PAR-CLIP data, which often contains low-complexity, partially degraded reads, or reads with mutations near protein-RNA interaction sites. To improve the alignment sensitivity, we adjust’ --winAnchorMultimapNmax 300’ and ‘ --seedSearchStartLmax 20 ‘, where the former increases the permitted number of mapping locations for each seed, aiding the alignment of repetitive regions that are common in non-coding RNAs. The latter reduces the maximum seed length that is used to start the alignment. Therefore, even in the absence of long exact matches of the seeds, STAR can initiate alignment. Collectively, these parameter settings enhance the ability to detect reads with weak or partial matches, making them specifically advantageous for CLIP-seq and PAR-CLIP data. All these parameters can be dynamically changed via the command-line parameters of ‘pcliptools-align’.

There are some additional parameters that were kept constant throughout the alignment procedure, i.e., ‘--alignSJoverhangMin 100 ‘ and ‘--alignSJDBoverhangMin 100 ‘, which set 100 nucleotides as the minimum overhang for unannotated and annotated junctions, respectively. These stringent thresholds reduce the probability of capturing false-positive splicing events by requiring substantial support at the splicing junction. To ensure reproducibility of alignment, the random number generator was fixed by using ‘-- runRNGseed 123’. Although these parameters were fixed in our pipeline, users have the ability to modify the source code if required.

### Peak calling by PCLIPtools

#### Calculate mismatch rate

Once the FASTQ file is aligned to the genome and a BAM file is generated, PCLIPtools performs peak calling by ‘pcliptools-cluster’. The first step involves extracting pileup information from the BAM file, which contains reference bases and sequences for all sequenced reads. This is accomplished via the ‘samtools mpileup’ function in the backend, which extracts the mismatch metrics essential for identifying RBA-RBP interaction sites.

For the ‘samtools mpileup’ function, the read depth cutoff is set to maximum to capture all genomic regions. However, to capture only high-quality reads, sequence mapping quality and base quality were set to 55 and 20, respectively (29). These two parameters can be changed via the command line depending on the data and users’ preferences. Since PAR-CLIP primarily induces T-to-C mismatches, we excluded insertion and deletion events in the mpileup output.

The mpileup function is performed in a strand-specific manner using only uniquely mapped reads to avoid ambiguity from regions that report bidirectional reads at a single genomic locus. To achieve this, uniquely mapped reads to either forward-strand or reverse-strand are captured via ‘ --excl-flags 272’ and ‘ --incl-flags 16 --excl-flags 256’ respectively. Custom bash functions embedded in the program eventually generate strand-specific bedGraph files that store the nucleotide-level positional information and respective mismatch count information. These bedGraph files are subsequently used to calculate the total number of bases sequenced and to calculate the non-T-to-C mismatch rate, a critical metric for identifying background noise in PAR-CLIP datasets.

#### Compute read groups and extract mismatch metrics

Contiguous regions in an alignment file with a minimum read depth are defined as read groups. These represent piles of reads that are putative interaction sites for RNA and RBP. To identify these regions, ‘samtools view’ in combination with ‘bedtools genomecov’ function are used. After applying minimum read depth filter, candidate read groups are extracted. To prevent fragmentation of the extracted read groups, neighboring regions are concatenated by ‘bedtools mergè.

We observed that there are very long stretches of reads in alignment files, which might represent true RNA-RBP interactions; however, in many cases, these could lead to false discovery, arising from reads allocated to repetitive elements, partial strand specificity, or mapping artifacts in low-complexity genomic regions. Such long stretches can artificially inflate the number of predicted binding sites and compromise the resolution of PAR-CLIP. To mitigate this, we split such regions longer than 50 bases into smaller read groups. Therefore, 50 is defined as the maximum width of the cluster, which can be changed from the command line interface while running the tool. Finally, the regions are saved as 6-column Browser Extensible Data (BED) file that contains location and strand information for each read group.

For all resulting read groups, the total number of bases sequenced, read depth, and number of T-to-C and non-T-to-C are quantified by intersecting read groups with previously generated strand-specific bedGraph files containing mismatch information.

#### Peak calling from read groups

PCLIPtools assume the following hypothesis for peak calling.

For whole transcriptome

Total base sequenced, 𝑁 = ∑ 𝐴𝑙𝑙 𝑠𝑒𝑞𝑢𝑒𝑛𝑐𝑒𝑑 𝑏𝑎𝑠𝑒𝑠

Total background mismatch, 𝑀 = ∑ 𝐵𝑎𝑐𝑘𝑔𝑟𝑜𝑢𝑛𝑑 𝑚𝑖𝑠𝑚𝑎𝑡𝑐ℎ𝑒𝑠

Transcriptome-wide normalized background mismatch rate is defined as 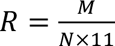

We assume there is a total of N read groups.

For 𝑖 ∈ {1, . . 𝑁}, let 𝑛_𝑖_denote the total sequenced base for the 𝑖^𝑡ℎ^ read group.

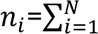 𝑏𝑎𝑠𝑒𝑠 𝑖𝑛 𝑖^𝑡ℎ^ read group

Expected number of T-to-C mismatches, 𝐸_𝑖_ = 𝑃𝑜𝑖𝑠𝑠𝑜𝑛(𝑅 · 𝑁)

When the observed number of T-to-C (𝑂_𝑖_) is higher than expected, the 𝑖^𝑡ℎ^ read groups are defined as a cluster.

The BED file containing the location and detailed metrics of the read groups is fed into an R script that performs statistical testing to identify high-confidence interaction sites. To do so, it performs a poison test,

poisson.test(O_i_,n_i_,R,alternative="greater")

and a p-value is obtained for each read group. Finally, False Discovery Rate (FDR) is calculated by the Benjamini-Hochberg (BH) method. Read groups with FDR ≤ 0.05 are considered a high-confidence interaction sites or clusters.

Afterwards, the respective clusters are annotated to the respective genomic regions.

#### Annotate clusters to genomic regions

For annotating the clusters, we developed a custom bash script by employing different functions from the ‘bedtools’ suite, ‘awk’, and ‘sed’. ENSEMBL (30) Gene Transfer Format (GTF) was used to allocate the clusters to corresponding genes. When a cluster overlapped multiple genomic features, the following hierarchy was used to resolve the annotation: 3’ untranslated regions (3’ UTR), coding sequence (CDS), 5’ untranslated regions (5’ UTR), and intron (16). If the associated gene’s biotype was non-protein-coding, then it was categorized with the appropriate gene category, such as miRNA, rRNA, etc. Clusters overlapping with multiple genes or multiple types of genes were flagged with a custom tag “multigenic” to flag annotation ambiguity. Point to be noted, as of PCLIPtools version 0.7.3, tRNAs are not assigned to the clusters due to their absence in canonical GTFs.

PCLIPtools also offers a stand-alone annotation script named ‘pcliptools-annotatè which allows users to annotate a standard 6-column BED file to the corresponding ENSEMBL GTF file. The output file retains the input 6 columns and additional 7^th^-9^th^ columns represent genomic features, i.e., 3’ UTR, 5’ UTR, etc, generic gene name, and ENSEMBL gene id, respectively.

### Alignment and Peak calling by Bowtie-PARalyzer pipeline

The GitHub repository named PARpipe (https://github.com/ohlerlab/PARpipe) is the wrapper for the Bowtie-PARalyzer pipeline. It employs Bowtie (v1.2.2) (31) as the aligner for sequencing, afterward PARalyzer and accompanying scripts in the PARpipe repository are used for peak calling and annotation.

### Processing, mapping of short-read data

The first two steps of the PARpipe pipeline perform read processing and alignment. Briefly, the ‘fastqToFASTA.pl’ script extracts the reads from the FASTQ file, and cutadapt removes the adapters and UMIs. Afterwards, the ‘collapseFA.pl’ script collapses the FASTA by removing duplicates. The resulting file is saved as the input FASTA file for Bowtie.

In the second step of PARpipe, Bowtie v1.2.2 is used to align the reads from FASTA files to the reference human genome version hg38. For the Bowtie, the following parameters are set ‘-v 2 -m 10 --best –stratà where -v 2 designates two mismatches are allowed per read and -m 10 discards any reads that have more than 10 valid alignments, thus reducing multi- mappers. The last two parameters allow Bowtie to report only the best alignments that have lower mismatches. The resulting BAM file is used by the rest of the steps to call peaks.

### Peak calling by PARalyzer

In the third step PARpipe generates a configuration file that contains the name of the input BAM file and retrieves the necessary parameters to run the PARalyzer. For all MATR3 PAR-CLIP, we used the default parameters of PARalyzer, some of which are MINIMUM_READ_COUNT_PER_GROUP=5, which keeps only the read groups with a minimum of 5 reads, MINIMUM_READ_COUNT_PER_CLUSTER=2 ensures the clusters or the predicted RBP-RNA interaction sites contain at least 2 reads, MINIMUM_READ_COUNT_FOR_KDE=3 for the kernel density estimation 3 reads are required, and MINIMUM_READ_COUNT_FOR_CLUSTER_INCLUSION=1. Although these defaults were used consistently, the order of priority logic (i.e., how overlapping conditions are handled internally by PARalyzer) was not explicitly documented and remains unclear.

Once PARalyzer completes peak calling, identifying the predicted T-to-C enriched RBP-RNA interaction sites, a series of perl scripts in the following PARpipe steps annotate the clusters to the genome by using a GENCODE (32) GTF v24 (https://ftp.ebi.ac.uk/pub/databases/gencode/Gencode_human/release_24/gencode.v24.a nnotation.gtf.gz) file.

Due to compatibility issues, PARalyzer could not process an updated GTF (ENSEMBL 112 or equivalent GENCODE GTF). To facilitate consistent downstream comparisons of the clusters resulting from different pipelines, all PARalyzer-derived clusters were reannotated to ENSEMBL GTF with ‘pcliptools-annotatè. This enabled direct comparison of the annotated genomic features across methods.

### Alignment and Peak calling by the STAR-PARalyzer pipeline

For the STAR-PARalyzer pipeline, we adopted the same strategy as used in the STAR- PCLIPtools workflow. For the MATR3 PAR-CLIP data, we directly used the BAM files generated by ‘pcliptools-align’. Additionally, for both DHX36 and HNRNPK, the same strategy and parameters were used as described for STAR-PCLIPtools alignment.

To perform peak calling with PARalyzer on STAR-generated BAM files, we wrote a custom script that mimics the peak calling steps of the PARpipe pipeline. For MATR3 PAR-CLIP data, we retained PARalyzer’s default parameters.

While comparing clusters resulting from PARalyzer and PCLIPtools, PARalyzer-specific MATR3 clusters showed low read depth compared to PCLIPtools-specific MATR3 clusters. Furthermore, the internal prioritization of PARalyzer’s read depth-related parameters was not explicit. To overcome this, we set all of the read depth parameters to 3, i.e., ‘MINIMUM_READ_COUNT_PER_GROUP=3, MINIMUM_READ_COUNT_PER_CLUSTER=3 MINIMUM_READ_COUNT_FOR_KDE=3, and MINIMUM_READ_COUNT_FOR_CLUSTER_INCLUSION=3’ for DXH36 and HNRNPK peak calling. The rest of the parameters were set to the default.

Once PARalyzer finished the peak calling steps, pcliptools-annotate was used to reannotate the PARalyzer-derived clusters with ENSEMBL GTF.

### Computing and plotting single nucleotide mismatches from BAM file

PCLIPtools offers a custom script ‘pcliptools-compute-substitution’ that computes all kinds of single nucleotide mismatches and internally generates a bar plot using ‘pcliptools- plot-substitution.R’. The former script accepts a BAM file and a genome FASTA file as minimal input. Additionally, a minimum read depth parameter can be provided that limits the read depth. By default, there is no limitation on the read depth parameter. Using the same mpileup module described in section ‘Calculate mismatch ratè, it computes the mismatches and forwards them to the latter script. The plotting script utilizes the base R function ‘`barplot()`’ to render the mismatches into a bar plot.

Another utility script ‘pcliptools-compute-bigwig’ is used to generate a bigWig file summarizing single-nucleotide mismatch count. Currently, it can compute Count Per Million (CPM) normalized bigWig for both T-to-C and non-T-to-C. The type of mismatch is determined by the parameter ‘-t’, which accepts either ‘T2C’ or ‘nonT2C’. It requires strand-specific bedGraph files generated by ‘pcliptools-cluster’ that store all the read depth and mismatch metrics (described in the section ‘Calculate mismatch ratè). Additionally, it requires a respective genome size file that contains the chromosome size information. This script extracts the requested mismatch information and writes it to a bedGraph file, which is converted to a bigwig file by the ‘bedGraphToBigWig’ function from the ‘UCSC kentUtils’ repository. Notably, the latter program is available as a standalone executable in https://hgdownload.soe.ucsc.edu/admin/exe/ and it does not need any additional installation process or admin privilege to execute.

### Comparison of cluster files via Venn diagram

All the Venn diagrams were generated using utility script ‘pcliptools-compare-bed’. This Python script relies on some additional libraries named argparse, pybedtools and matplotlib-venn which are not part of the core PCLIPtools peak calling pipeline. This script accepts a maximum of three BED files to render a proportional Venn diagram. The BED files are read using pybedtools library. Afterwards, comparison of the coordinates is performed by ‘bedtools multiinter’. A matrix is generated that contains the overlap information. Depending on the number of datasets, ‘venn2()’ or ‘venn3()’ function from ‘matplotlib- venn’ library plots the Venn diagrams and saves the image as files.

If the cluster BED files contain the target gene names in one of the columns, it can compare gene-level overlap instead of the cluster coordinates. Since the script uses ‘matplotlib- venn’ library which can handle maximum of three data sets to generate Venn diagram, this is the highest limit for pcliptools-compare-bed.

### Correlation of clusters by T-to-C mismatches

Correlation analysis of clusters was performed by computing the total number of T-to-C mismatches per gene. Then, the R function `cor()` was used to calculate the correlation matrix. The R library ‘pheatmap’ was used to plot the heatmap illustrating correlation (**Fig. 2D**).

**Figure 2.**
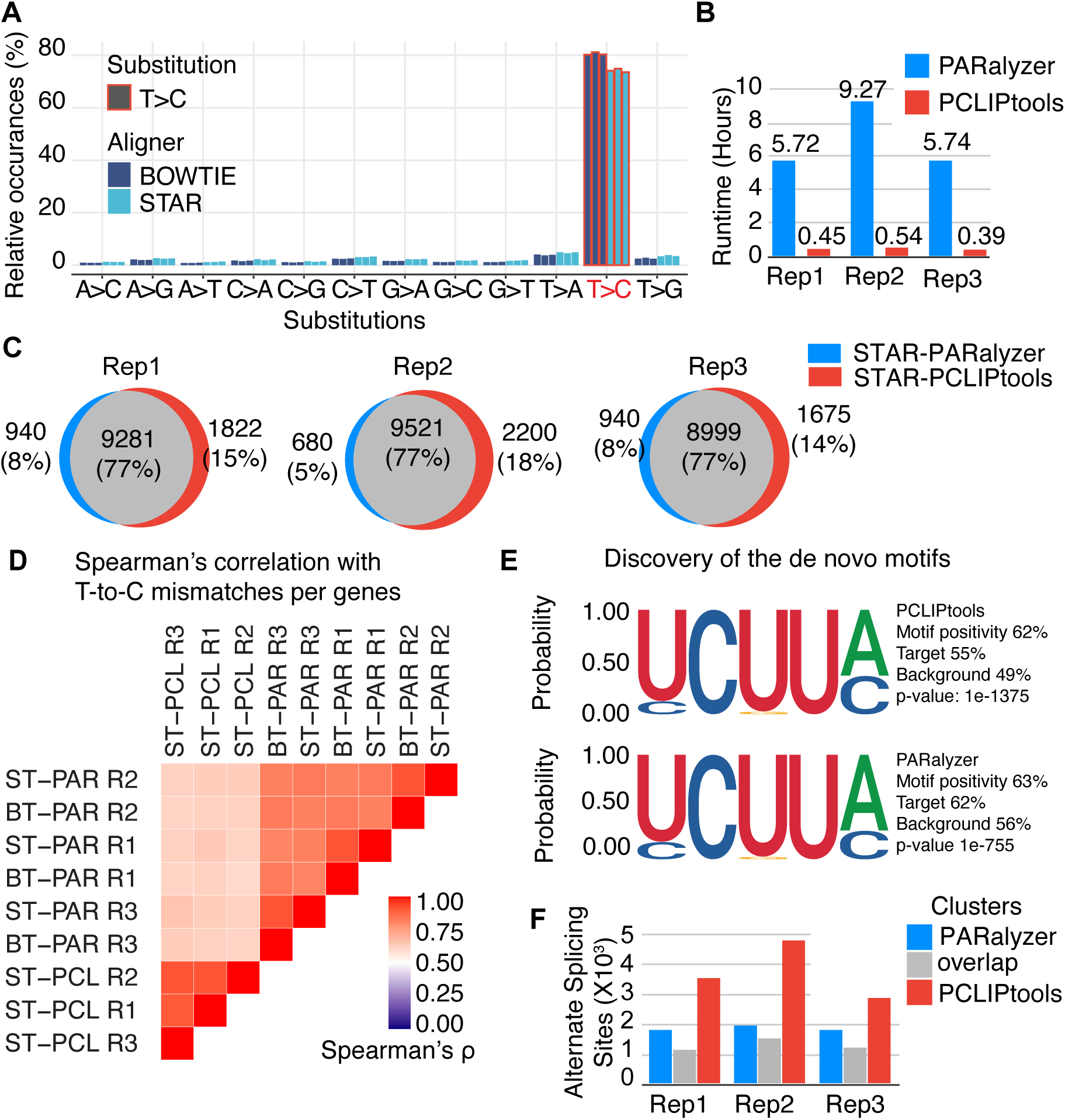
(**A**) PAR-CLIP shows characteristic T-to-C mismatches (T>C) regardless the aligner used. (**B**) Usage of unified BAM files (STAR generated) resulted in higher speed for PCLIPtools compared to PARalyzer. (**C**) PCLIPtools derived targets are comparable to PARalyzer derived targets and (**E**) result in similar binding motif, while additionally identifying novel targets not detected by PARalyzer. (**D**) STAR-PCLIPtools resulted in highly reproducible clusters accross replicates compared to Bowtie-PARalyzer or STAR-PARalyzer, ST : STAR, BT : Bowtie, PAR : PARalyzer, PCL : PCLIPtools; R1, R2, R3 depicts three replicates. (**F**) PCLIPtools specific clusters are functionally more associated with AS events.

### Motif discovery

Homer (33) version 4.10 was used to compute the *de novo* motif. The resulting cluster coordinates were provided to findMotifsGenome.pl function of Homer, and the length parameters for the k-mer were set to 5,8, and 10.

### Read depth calculation for clusters

Mean read depth calculation was performed using ‘plotCoveragè function from deepTools v3.5. This function accepts BAM files and cluster BED as input and provides two informative plots. One plot shows the mean read depth for the clusters vs the fraction of clusters that contain the reads. The second plot is a cumulative plot that shows read coverage vs fraction of reads that are higher than the read (**Supplementary Fig. S4**). This function also output the raw counts of reads for each cluster that can be used to plot mean read depth per sample (**Fig. 3A**).

**Figure 3.**
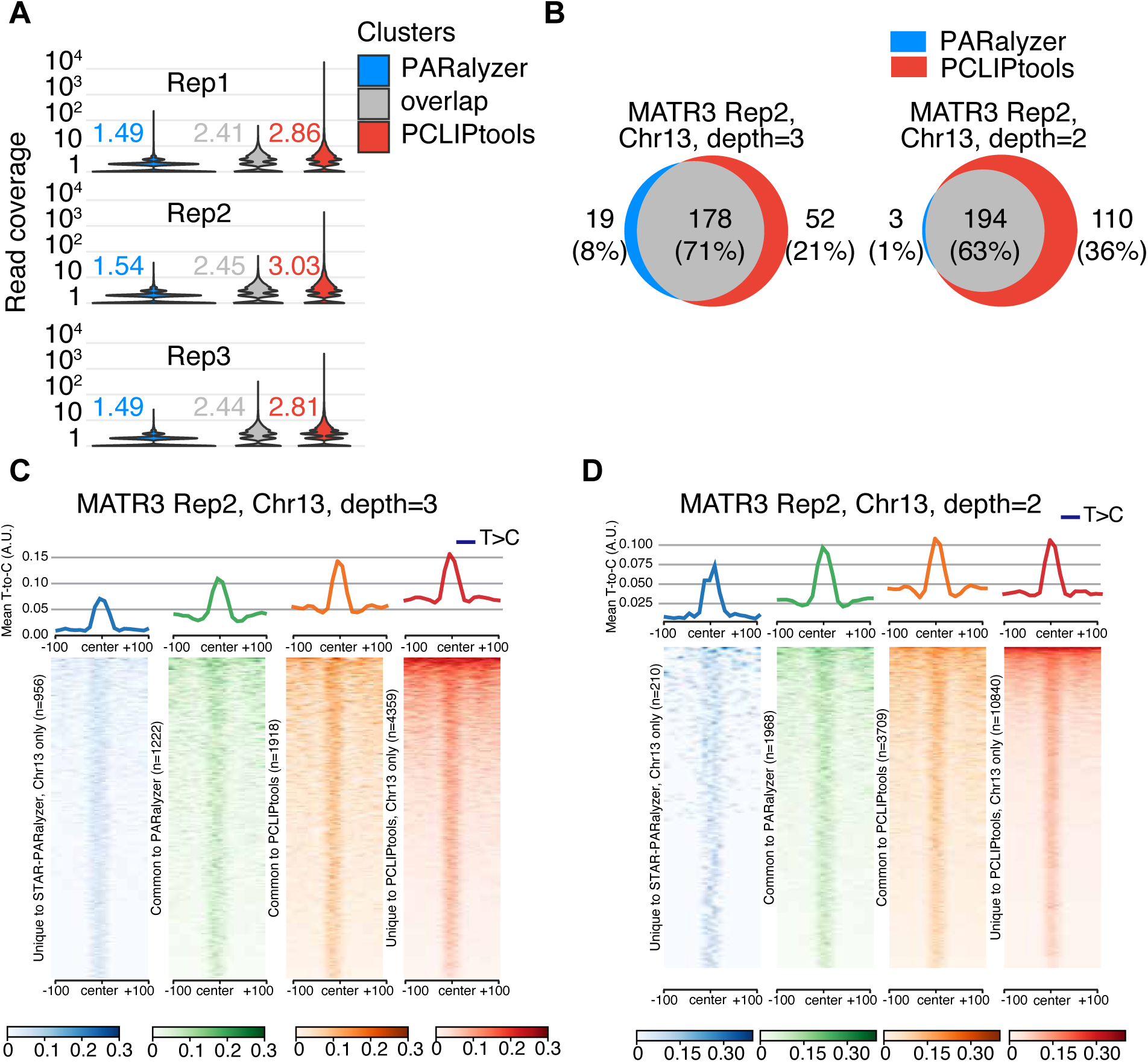
(**A**) Transcriptome-wide PCLIPtools specific clusters are higher in read depth compared to PARalyzer specific clusters. (**B**) Chr13 only analyses showed PCLIPtools started capturing PARalyzer specific clusters upon reducing minimum read depth cutoff. (**C**) For both read depth 3 and (**D**) read depth 2 threshold PCLIPtools identify clusters abundant with T-to-C conversion events whereas PARalyzer fails to detect those high confidence sites.

### Comparison of T-to-C mismatch across clusters

Comparison of the T-to-C mismatch enrichment across different clusters was performed via deepTools. First, a bigWig file containing T-to-C mismatches was generated by ‘pcliptools- compute-bigwig’. The resulting bigWig files and the cluster BED files were provided to ‘comoputeMatrix referencè function of deepTools. It produced matrices of T-to-C densities across clusters. Finally, ‘plotHeatmap’ or ‘plotProfilè of ‘deepTools’ was used to generate the heatmap and line plot to visualize T-to-C enrichment over the cluster regions.

### Comparing BAM file for similarity

To evaluate whether multiple alignment files generated comparable alignment, the BAM files were compared. To do so, ‘multiBamSummary’ from ‘deepTools’ was used. The program divides the genome into equal-sized genomic bins and then calculates the number of reads across those bins for all BAM files. Afterwards, the read number per genomic bin is used to compute Spearman’s correlation, and a matrix is generated. Finally, ‘plotCorrelation’ of ‘deepTools’ is used to plot a correlation heatmap from the resulting matrix.

### Hardware Configuration

The NIAMS HPC cluster was used for all computations. Job submissions were managed via SLURM batch scripts, which allocated 16 cores and 32 GB of memory for each job. GNU Parallel (34) was used to parallelize the queues.

## Results

### PCLIPtools uses a simple methodology to call peaks

PCLIPtools is divided into two basic modules; the first one performs alignment of the sequencing data, and the second one performs peak calling. The aligner module first collapses the reads from FASTQ files to remove PCR duplicates. The adapters and unique molecular identifiers (UMI) are removed by utilizing cutadapt (v4.9) (28). The resulting FASTA files are aligned to the desired genome by STAR (23) to obtain a Binary Alignment Map (BAM) file (**Fig. 1B**). Finally, this alignment file is used for peak-calling. PCLIPtools extracts all the transcriptome-wide mismatch events and categorizes T-to-C as signal and the rest as background (**Fig. 1C**). Upon calculating the rate of transcriptome-wide background mismatches, a Poisson model is computed. Among the aligned reads, contiguous reads in a genomic region that surpass a certain read depth are considered as read groups. The Poisson model is used to calculate the expected number of T-to-C mismatches, and the read groups having more T-to-C mismatches than the expected rate are considered high-confidence RNA-RBP interaction sites. Finally, the confident sites are annotated to genomic regions with help of a reference General Feature Format (GTF) file.

### PCLIPtools outperforms PARalyzer in runtime efficiency and higher cluster detection

We decided to benchmark PCLIPtools against a single PAR-CLIP analysis software, PARalyzer, for two reasons: 1) PARalyzer is widely adopted for PAR-CLIP (**Fig. 1A**) and 2) we experienced installation or execution issues with wavClustR and BMix. As test datasets, we settled on a recent set of experiments identifying the binding sites of MATR3 (25), because: 1) MATR3 is an abundant nuclear protein binding to a wide-range of exonic and intronic sequences (35), 2) the experiments were performed in biological triplicates, and 3) the PAR-CLIP libraries were sequenced deeply with 35M reads per sample, allowing us to challenge computational resources.

We performed data pre-processing on three replicates of MATR3 PAR-CLIP data from the HAP1 cell line (25). FASTQ files were first collapsed into FASTA format by removing duplicate reads that resulted in 13, 17, and 13 M reads for the three replicates (**Supplementary Table 1**). Next, adapters and unique molecular identifiers (UMIs) were trimmed by using cutadapt.

More than 91% reads had adapters (**Supplementary Table 1**). The obtained processed sequence reads were aligned to the human genome version hg38 with STAR (2.7.9a) for PCLIPtools. We chose STAR as the preferred aligner because of its speed, accuracy, and ability to map spliced reads (23). PARalyzer is wrapped under a pipeline named PARpipe (https://github.com/ohlerlab/PARpipe) (36) that uses Bowtie (31) for alignment. Both aligners yielded 6-8 M unique mappers (**Supplementary Table 1**). The resulting BAM files were subsequently used as input for peak calling.

PARalyzer initiates a Java program that processes BAM files to identify genome-wide contiguous regions (read groups) mapped by NGS reads. For each of the read groups, it estimates a signal kernel density based on T-to-C mismatches and a background kernel density based on T-to-T matches. Regions with a higher signal-to-background likelihood are classified as interacting regions or clusters (16). For the three replicates of MATR3 PAR-CLIP PARalyzer identified 116,661, 114,794, and 109,165 clusters (**Supplementary Table 2**) upon running for a mean of 10.5 hours. PCLIPtools calls peaks by modeling the Poisson distribution of background mismatches and then identifying regions where T-to-C mismatches are significantly higher than expected under the background model. For the three replicates, PCLIPtools identified 246,270, 319,431, and 193,332 clusters (**Supplementary Table 2A**) that passed statistical testing for the same replicates upon running for mean 0.5 hours (**Fig. 1D**). There were 21,582, 25,927, and 19,868 read groups that failed the hypothesis tests as their FDR was greater than the threshold of 0.05 and thus were not considered as confident clusters. **Table 2A** summarizes the resulting cluster numbers.

The obtained clusters were annotated with ENSEMBL (30) derived GTF (v112) (https://ftp.ensembl.org/pub/release-112/gtf/homo_sapiens/Homo_sapiens.GRCh38.112.chr.gtf.gz) using the annotation script designed for PCLIPtools to categorize the transcript regions from which they were derived. Regardless of the varying total number of clusters across all replicates and tools used, introns were, as expected, the main enriched interaction region for MATR3 (**Fig. 1E**). Roughly 80% of the clusters aligned to intronic sequence space, and 3’UTR accounted for another 8-10% of the clusters. Collectively, these results suggest that STAR-PCLIPtools required only a fraction of the time needed by Bowtie-PARalyzer for alignment and peak calling while detecting a substantially higher number of peaks.

### The number of resulting clusters is affected by the aligners and the peak calling method

One obvious rationale for the higher number of clusters for PCLIPtools compared to PARalyzer is the difference in strategy that is used to break down the read groups into smaller chunks of genomic regions. Although there is a minimum width parameter (stretch of nucleotides to make a read group) for the clusters in PARalyzer, there is no threshold for maximum width, often resulting in multiple independent, but close sites being counted as a single, implausibly long cluster, considering that most RBPs will have a footprint that is <20 nucleotides (nt). To avoid this phenomenon, we set the maximum width of the cluster to 50 nt by default (of course, further changeable via command line parameters). As a result, read groups more than 50 nt larger are broken down into multiple read groups, and a statistical test is performed for all of them. By doing so, only smaller read groups with high confidence will be reported as clusters. Even after merging the adjacent clusters from PCLIPtools to a single cluster, it retains 224,191, 282,290, and 178,215 clusters for three replicates, resulting in a higher number of clusters for PCLIPtools (**Supplementary Table 2A**, column STAR- PCLIPtools-merged). Therefore, the breakdown strategy of the read groups by our method is not a major source of high cluster yield for PCLIPtools.

Considering that we used a different genomic aligner to obtain a vastly increased number of clusters using PCLIPtools compared to PARalyzer, we wondered whether the aligner itself could account for the difference in identified clusters. The number of reads aligned with both STAR and Bowtie for all three replicates was comparable (7,8 and 6 M, respectively, for replicates 1, 2, and 3; **Supplementary Table 1**). However, the read distribution across the genome was not identical. A similarity analysis using ‘multiBamSummary’ of deepTools (37) (version 3.5), quantified and compared multiple alignment files with a 10,000 base sliding window, revealed that Bowtie and STAR-based alignments were similar, with Spearman’s correlation ranging from 0.72 to 0.76 (**Supplementary Fig. S1**).

Both aligners captured a similar pattern of T-to-C mismatches – the diagnostic mutation introduced by crosslinking - showing a higher frequency of T-to-C conversions compared to other mismatch types (**Fig. 2A**). However, 80% of the mismatches were T-to-C for Bowtie, whereas STAR counted 75% mismatches as T-to-C. For analyzing PAR-CLIP data, read depth is a critical factor in downstream processing, with higher read coverage providing greater confidence in results (38). Taken together, we did not observe appreciable differences in read depth or alignment preferences between the two aligners (**Supplementary Table 1)**.

STAR is considered a super-fast aligner and is able to align 309 million read pairs/hour with six computational cores (23), whereas Bowtie aligns only 88.1 million reads/hour with four cores (31). This indicates that at least part of the runtime differences are likely resulting from the choice of aligner. Therefore, we adapted PARalyzer to use the STAR-generated alignment as input to limit peak callers to a single variable, allowing for a more accurate comparison of resulting cluster counts and computation times. Despite using the same input file, PCLIPtools once again outperformed PARalyzer in computation time (**Fig. 2B**). Across three replicates, PARalyzer required an average of 6.91 hours for peak calling, whereas PCLIPtools completed the task in an average time of 0.46 hours. This underscores the significant reduction in peak calling time achieved by PCLIPtools. Notably, there was a slight decrease in the number of clusters PARalyzer called using STAR rather than Bowtie alignments as input. Across three replicates, STAR-PARalyzer resulted in 99,551, 99,850, and 95,240 clusters (**Fig. 1E, Supplementary Table 2)**. The lower yield for STAR-PARalyzer-derived clusters may be due to a slightly reduced number of T-to-C containing reads captured by STAR (**Fig. 2A**). Nevertheless, even though cluster numbers differed, the same proportion (82%) of the clusters aligned to intronic regions (**Fig. 1E)**. Taken together, our benchmarking indicates that PCLIPtools offers significant advantages over PARalyzer with reduced runtime and higher peak numbers. In the subsequent sections, we explored whether PCLIPtools- derived clusters are biologically comparable with PARalyzer-derived clusters.

### PCLIPtools and PARalyzer clusters show high concordance for transcripts of origin

Although the majority of clusters called by PARalyzer and PCLIPtools were aligned to intronic regions, we sought to understand whether they are similar and if the predicted gene regulatory outcomes are comparable to each other. To do so, we checked the reproducibility of the clusters by performing intra-tool and inter-tool cluster comparisons. First, we compared the genomic coordinates of the Bowtie-PARalyzer and STAR-PCLIPtools derived clusters, and our initial finding suggests that for each of the replicates, approximately 20% of the union of the clusters overlaps between Bowtie-PARalyzer and STAR-PCLIPtools (**Supplementary Fig. S2A and Supplementary Table 2**). There was a 41-48% union of the clusters that were unique to PCLIPtools and were not detected by PARalyzer. Similarly, 32-39% clusters were unique to PARalyzer that were not identified by PCLIPtools. Nonetheless, the two approaches captured largely the same target gene set, with a 76–77% overlap in the union of the two sets (**Supplementary Fig. S2B and Supplementary Table 2**). When using PCLIPtools and PARalyzer clusters derived from the same BAM files, i.e., STAR-PARalyzer and STAR-PCLIPtools, we observed a similar trend. Across all three replicates, approximately 19-20% of the union of the clusters overlaps between STAR-PARalyzer and STAR-PCLIPtools (**Supplementary Fig. S2C and Supplementary Table 2**). We also found that 42-49% of clusters were unique to PCLIPtools and were not detected by PARalyzer. Additionally, 32-38% of clusters were unique to PARalyzer and were not detected by PCLIPtools (**Supplementary Fig. S2C and Supplementary Table 2**). Again, a larger (77%) set of target genes was captured by both tools (**Fig. 2B**), suggesting more similarities in the results when comparing target gene levels but substantial dissimilarities in identified clusters.

To address the discrepancies in the cluster coordinates, we first assessed the reproducibility of clusters from different biological replicates of MATR3 PAR-CLIP using the same tool. The union of the clusters from Bowtie-PARalyzer and STAR-PARalyzer showed a 2% and 3% overlap, respectively, while those from STAR-PCLIPtools clusters exhibited a more significant overlap of 11%. This 11% consisted of 60,322 clusters shared across all three replicates (**Supplementary Fig. S3**), suggesting that PCLIPtools yields a more reproducible outcome. It is important to note that the low overall reproducibility of clusters across biological replicates likely arises from the fact that the PAR-CLIP experiments were not saturating, meaning they did not capture all possible MATR3 binding sites defined by a short sequence element that frequently occurs in the vast intronic space co-transcriptionally occupied by MATR3 (35). Consistently, MATR3 PAR-CLIP replicates captured largely the same set of transcripts (**Supplementary Fig. S3**), whether identified by PARalyzer or PCLIPtools. The similarity matrix of the BAM files computed via deepTools revealed that alignments by Bowtie and STAR were highly comparable, showing Spearman’s correlation ranging from 0.72 to 0.76 (**Supplementary Fig. S1**). However, inter-replicate comparisons indicated that STAR created slightly more uniform alignments compared to Bowtie, showing Spearman’s correlations of 0.61-0.64 and 0.56-0.59, respectively. This further suggests that the choice of aligner underlies the emergence of dissimilar clusters. Finally, calculating similarity of the PAR-CLIP datasets by quantifying T-to-C conversion numbers per transcript showed that the replicates analyzed by PCLIPtools showed a very high correlation (Spearman’s ρ ∼0.9), slightly higher than PARalyzer (Spearman’s ρ ∼0.83) (**Fig. 2D**). Taken together, PCLIPtools not only outperforms PARalyzer in runtime but also produces relatively higher degrees of reproducible results across replicates.

### Clusters called by PCLIPtools contain the canonical MATR3 binding motif

Considering the differences in clusters identified by PARalyzer and PCLIPtools, we next sought to examine the source of these discrepancies and whether one or both tools exhibited a substantial false positive rate for binding site determination. Most RBPs interact with RNA at typically short and clearly defined sequence or structural elements; in the case of MATR3, these include the UCUU[AC] in HepG2 (39) and HCT116 cells (35). UCUU[AC] and CCGUA in the HAP1 cell line (25). *In vitro*, in RNAcompete assays, MATR3 bound AUCUU (40). Therefore, we investigated the enrichment of sequence elements in our PARalyzer and PCLIPtools clusters, under the assumption that the MATR3 binding motifs should be enriched at true positive interaction sites. After combining tool-specific clusters from all three replicates of MATR3, we computed a de novo motif with Homer (ref + version) and observed an enrichment of the UCUU[AC] motif (**Fig. 2E**). Out of 218,998 combined clusters from PARalyzer, 139,487 (63%) contained the UCUU[AC] motif. On the other hand, of the 364,705 combined clusters from PCLIPtools, 226,927 (62%) contained the motif. This confirms that both tools are capturing the same signals, but PCLIPtools demonstrates greater robustness and homogeneity in motif discovery as the UCUU[AC] motif is found at 49% background sequences compared to 56% for PARalyzer. We also identified clusters unique to each tool and computer *de novo* motif. The result showed a similar motif and indicated that these were true binding sites (data not shown), and the reason they were not captured by the other tool was not due to being noise.

Next, we assessed whether the PCLIPtools clusters also enrich functional binding sites, defined as binding sites present on transcripts that are affected by the depletion or overexpression of the studied RBP. Previously, we found that MATR3 knockout (KO) leads to changes in the alternative splicing (AS) of transcripts with MATR3 binding sites near the 5’ and 3’ splice sites (25,35). By integrating the AS data from MATR3 KO in HAP1 cells (25) with our PAR-CLIP data, we discovered that a significant number of AS events co-localize with clusters identified by PCLIPtools and PARalyzer. Notably, clusters specific to PCLIPtools exhibited a greater enrichment of AS events compared to those unique to PARalyzer (**Fig. 2F**). These findings indicate that PCLIPtools captures biologically relevant MATR3-RNA interaction sites more effectively, including unique regions with potential functional significance.

### PCLIPtools deprioritizes low-coverage regions for peak calling

Next, we attempted to understand why PCLIPtools and PARalyzer each identified a number of unique clusters. We began by examining the read-depth and mismatch counts for tool-specific clusters. To minimize sampling bias due to population size, we conducted bootstrapped sampling of 10,000 regions from each category for each replicate and computed the mean read depth. Our comparison showed that the mean read-depth for PARalyzer-specific clusters was approximately 1.49 reads, compared to 2.41 reads for common clusters and 2.86 reads for PCLIPtools-specific clusters (**Fig. 3A**). The average read-depth for PARalyzer-specific clusters was almost 2 times lower than that of common or PCLIPtools-specific clusters, indicating that PCLIPtools rejected calling peaks in regions with low read-depth following alternative hypothesis testing.

To explore differences in tool-specific clusters, we limited the analysis to a small, manageable subset. We restricted the alignment file from replicate 2, due to its highest read depth (see **Supplementary Table 1**), to reads from chromosome 13 (chr13), which includes the gene *TDRD3*, identified as a key target of MATR3 (25), for further analysis.

On chr13 of replicate 2, we found 6,277 and 2,178 clusters for PCLIPtools and PARalyzer, respectively. While comparing target genes, 178 genes comprising 71% of the union of target genes were common for each tool. Fifty-two genes were unique to PCLIPtools, and 19 genes were unique to PARalyzer (**Fig. 3B**). However, while comparing the genomic coordinates, 19% of the union of the clusters were shared. Among the 6,277 clusters identified by PCLIPtools and the 2,178 clusters identified by PARalyzer, 1,918 and 1,222 clusters, respectively, overlapped between the two tools. We designated them as common-PCLIPtools and common-PARalyzer, respectively. These left 4,359 out of 6,277 PCLIPtools clusters and 956 out of 2,178 PARalyzer clusters unique to each tool. Computing the read density around these regions revealed that 4,359 PCLIPtools-specific clusters had approximately three times higher read density with a mean of 5.8 reads (**Supplementary Fig. S4A**) compared to 956 PARalyzer-specific clusters with a mean read depth of 1.8 (**Supplementary Fig. S4B**). As expected, T-to-C conversion events were also enriched in PCLIPtools-specific regions with a maximum value of 0.15 A.U. compared to those of PARalyzer, with 0.07 A.U., respectively (**Fig. 3C**). Similar read enrichment and T-to-C conversion were also observed for common-PCLIPtools over common-PARalyzer clusters. This indicated that PARalyzer not only missed certain high-confidence (high-read, high-conversion) interaction sites, but also identified low-read, low-conversion regions that are likely false positives. Taken together, our analyses reinforce our previous observations that PCLIPtools assigns low confidence to regions with low read depth and low T-to-C mutation signature, consequently discarding them as potential interaction sites.

### Relaxing PCLIPtools read-depth parameters results in a complete representation of PARalyzer clusters

Under default conditions, PARalyzer requires a minimum read depth of 5 reads for read groups, two reads per cluster, and three reads for inclusion into kernel density estimation. Thus, clusters called by PARalyzer should be expected to contain at least three reads. However, 50% of the unique PARalyzer clusters showed read depth lower than two reads (**Supplementary Fig. S4B**), likely due to errors in the hierarchy controlling the selection of read group and cluster. For PCLIPtools, we initially set three reads as the minimum requirement for a cluster, which is well reflected in the coverage plot of unique PCLIPtools clusters, as 100% of the clusters contained a minimum of three reads (**Supplementary Fig. S4A**). As noted above, PCLIPtools missed several PARalyzer-specific clusters with low read depth. Therefore, we tested whether relaxing the read depth parameter for PCLIPtools could capture additional clusters previously identified only by PARalyzer. Indeed, upon changing the minimum read-depth parameter from 3 to 2 for PCLIPtools, we found a total of 14,549 clusters on chr13 for MATR3 rep2. Upon reducing the read depth to 2, comparison of the target genes yielded 194 as a common target that comprised 63% of the union of genes and only 1% of the union of the genes remain unique to PARalyzer that were not captured by PCLIPtools (**F**i**g. 3B**). This suggests that reducing the read depth allows PCLIPtools to capture the previously ignored targets. Further analysis showed that of 14,549 clusters from PCLIPtools, 10,840 were unique to the tool and 3,709 had overlap with 1,968 clusters from PARalyzer. PARalyzer-specific cluster showed a drastic drop in number from 956 (**Fig. 3C**) to 210 (**Fig. 3D**). Again, PCLIPtools-specific clusters exhibited higher read depth with a mean of 3.9 reads (**Supplementary Fig. S4C**), compared to PARalyzer-specific clusters having a mean of 1.7 reads (**Supplementary Fig. S4D**). Additionally, 100% of the PCLIPtools-specific clusters contained a minimum of 2 reads, whereas 60% of the PARalyzer-specific clusters contained fewer than two reads. Upon comparing T-to-C enrichment, PCLIPtools specific 10,840 clusters showed higher conversion events with a maximum value of 0.1 A.U. compared to 210 PARalyzer specific clusters showing a maximum of 0.075 A.U. (**Fig. 3D**). Thus, relaxing the parameters of PCLIPtools allowed us to capture the low confidence clusters that were unique to PARalyzer and were initially set aside as not being robust enough for our peak calling algorithm.

### Reanalysis of DHX36 and HNRNPK PAR-CLIP data confirms the robustness of PCLIPtools peak calling

Next, we wanted to demonstrate that PCLIPtools provides robust results independent of the RBP studied by PAR-CLIP. We downloaded publicly available PAR-CLIP data for two additional RBPs, DEAH-box helicase 36 (DHX36) (26) and Heterogeneous nuclear ribonucleoprotein K (HNRNPK) (27), and used PCLIPtools to analyze the data. DHX36 is a cytoplasmic helicase that binds mRNA, and its role in unwinding G-quadruplexes is well studied. Sauer and colleagues (26) showed that DHX36 preferentially binds coding regions and 3’ UTRs of mRNAs. HNRNPK participates in pre-mRNA processing in the nucleus. Fallatah and colleagues (27) showed the role of HNRNPK in the shuttling of the target mRNAs to the cytoplasm.

Using the same alignment strategy as described above, we aligned the sequencing data with the STAR aligner and performed peak calling with both PARalyzer and PCLIPtools. A runtime comparison showed that for both DHX36 and HNRNPK, PCLIPtools was able to finish peak calling within 7 and 13 minutes, respectively. In contrast, PARalyzer took 115 and 125 minutes, respectively (**Fig. 4A**). For DHX36, PARalyzer found 13,759 clusters, whereas PCLIPtools identified 25,042 clusters (**Fig. 4B**). As expected, the major annotation for DHX36 was found to be 3’UTR and coding regions for both tools. For HNRNPK, PARalyzer identified 46,360 clusters, and PCLIPtools identified 105,231 clusters. The major annotations for both tools were introns and 3’UTR, in line with the published findings (**Fig. 4B**). Taken together for both DHX36 and HNRNPK PAR-CLIP PCLIPtools identified substantially higher numbers of clusters with retaining annotations from previously published data, and PCLIPtools took significantly lower runtime compared to PARalyzer.

**Figure 4.**
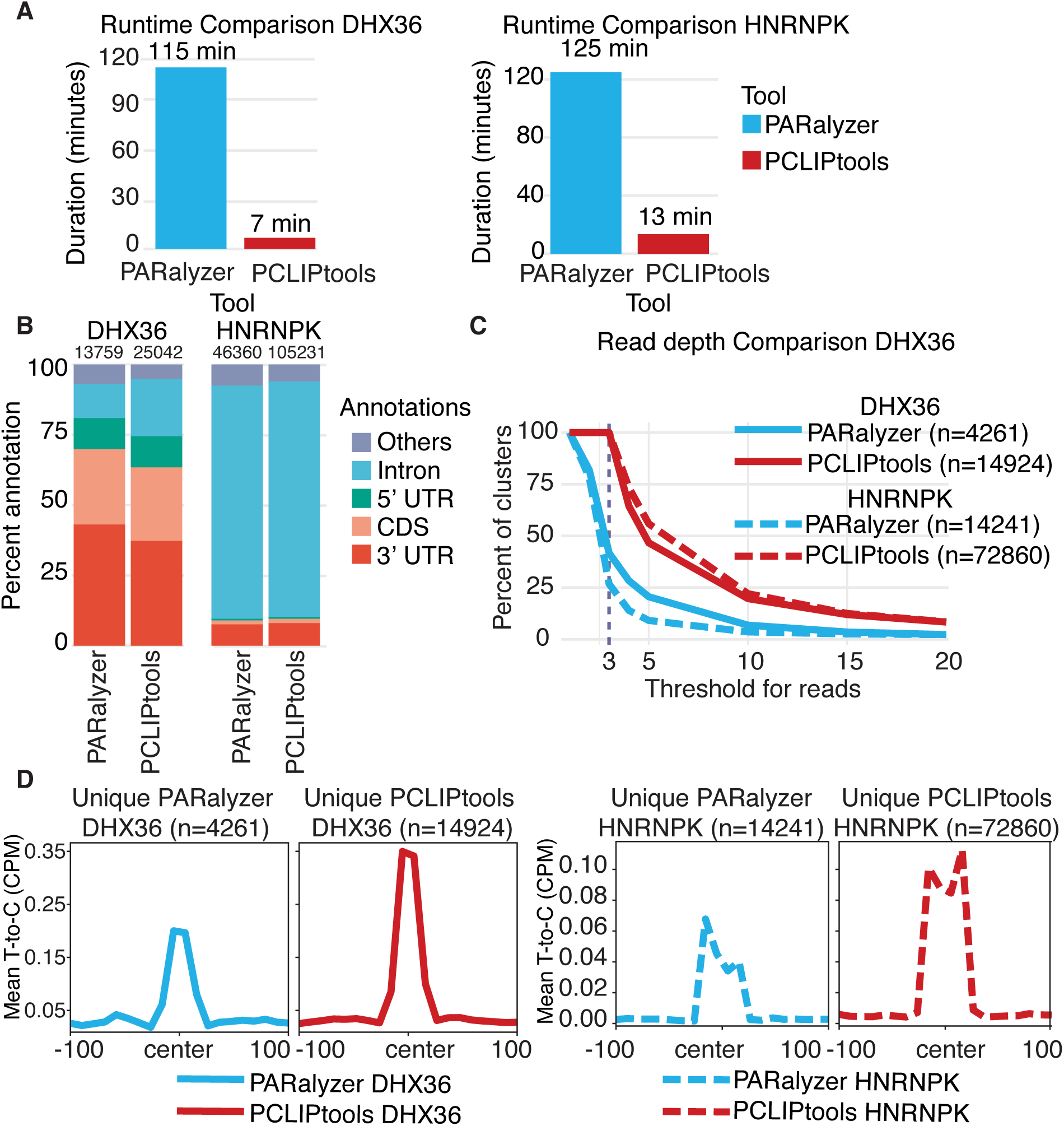
(**A**) Publicly available DHX36 and HNRNPK PAR-CLIP data shows PCLIPtools takes less runtime compared to PARalyzer but (**B**) yield substantially higher number of clusters with retaining primary annotations. Read depth and T-to-C mismatch distribution for tool specific unique clusters showed (**C**) PCLIPtools specific clusters are enriched with more reads and (**D**) higher T-to-C mismatch compared to PARalyzer specific clusters. UTR, untranslated regions.

For both DHX36 and HNRNPK PAR-CLIP, we identified tool-specific clusters that were identified by either of the tools. We found 4,261 and 14,241 clusters that were unique to PARalyzer for DHX36 and HNRNPK, respectively. 14,942 and 72,860 clusters were exclusive to PCLIPtools for DHX36 and HNRNPK, respectively (**Figs 4C and 4D**). Similar to MATR3, for both DHX36 and HNRNPK we found higher number of PCLIPtools-specific clusters than PARalyzer-specific clusters.

We wanted to evaluate if there were read depth and T-to-C conversion discrepancies across the tool-specific clusters. Read depth distribution for all tool-specific clusters showed PCLIPtools-specific clusters are high in read depth for both DHX36 and HNRNPK (**Fig. 4C**). For both PAR-CLIP datasets, all of the PCLIPtools-specific clusters contained at least three reads, which was in accordance with the set parameter. For PARalyzer, we exclusively modified the following parameters: ‘MINIMUM_READ_COUNT_PER_GROUP=3, MINIMUM_READ_COUNT_PER_CLUSTER=3, MINIMUM_READ_COUNT_FOR_KDE=3, MINIMUM_READ_COUNT_FOR_CLUSTER_INCLUSION=3’ to make sure the minimum read depth threshold for cluster is strictly set to three. However, despite setting the minimum read threshold as three, more than 60% of PARalyzer-specific DHX36 clusters contained fewer than three reads, and 75% of PARalyzer-specific HNRNPK clusters contained fewer than three reads (**Fig. 4C**). These demonstrate PARalyzer calling clusters in low read depth regions, ignoring the minimum low read depth parameter. PCLIPtools specific clusters for both PAR-CLIP also showed high enrichment of T-to-C conversion, with a maximum of 0.35 Counts Per Million (CPM) for DHX36 and 1.1 CPM for HNRNPK. PARalyzer-specific clusters, on the other hand, showed 0.20 CPM for DHX36 and 0.07 CPM for HNRNPK (**Fig. 4D**). Taken together, both PAR-CLIP and the PCLIPtools specific clusters were enriched with read depth and T-to-C mismatch, and thus increasing the confidence in the results. Thus, clusters missed by PCLIPtools can be safely disregarded.

Collectively, we demonstrated that PCLIPtools deprioritized low read-depth and low conversion-event regions for peak-calling, explaining the discrepancies between PARalyzer and PCLIPtools. Nevertheless, despite maintaining stricter default parameters, PCLIPtools was able to identify more interaction sites for MATR3 that captured the appropriate signals and were functionally relevant.

## Discussion

PAR-CLIP is an effective method for identifying RBP-RNA interactions that facilitates study of different RBPs. However, without an appropriate, easy-to-use tool, data analysis often becomes cumbersome for the biologists. The development of PCLIPtools had two things in mind: being robust in statistics and becoming user-friendly. Hence, Bash was the choice of programming language, so that most of the biologist-turned computational biologists can run the tool and can modify it as per their requirements. Instead of writing new programs or libraries, we utilized or repurposed built-in features of industry-standard stable tools, such as samtools (41) and bedtools (42). Any given high-performance computer used by computational biologists typically has those pre-installed. Therefore, we eliminated a frequent source of problems related to the installation of specialized software. Additionally, in our comparative analysis, it has been shown that PCLIPtools is superior to the widely adopted PARalyzer both in speed and cluster discovery. PCLIPtools-derived clusters were not only rich in reads but also had a high abundance of T-to-C conversions, ensuring the authenticity of the clusters. Correlation analysis of the PCLIPtools-derived clusters across replicates showed higher reproducibility. Most importantly, a higher number of PCLIPtools-derived clusters were positive for the consensus motif and confirmed the true nature of the RBP interactions. Comparison of the functional relevance with alternate splicing also confirmed higher performance of PCLIPtools-derived clusters.

A potential challenge for PCLIPtools was its inability to call the ∼5-8% of targets that were PARalyzer-specific. Our extensive follow-up computations showed PCLIPtools left those PARalyzer-specific clusters due to the low read and low T-to-C event counts and could be captured by relaxing mapping parameters in PCLIPtools. The low read depth indicates that these clusters are not highly occupied by RBPs and are unlikely to act as cis-acting elements in the regulation of gene expression, and will be of low priority for follow-up study. While working with NGS data, it is crucial for the users to set appropriate parameters to reduce noise and minimize the risk of false discovery. We believe PCLIPtools sets a new standard for PAR-CLIP data analysis to provide such high confidence backed by high read depth and high conversion events.

## Acknowledgements

This research was supported by the Intramural Research Program of the National Institutes of Health (NIH). The contributions of the NIH authors are considered Works of the United States Government. The findings and conclusions presented in this paper are those of the authors and do not necessarily reflect the views of the NIH or the U.S. Department of Health and Human Services. This work was supported by the Intramural Research Program of the NIH, National Institute of Arthritis and Musculoskeletal and Skin Diseases (ZIA-AR041205 to M.H.).

We would like to thank members of Astrid Haase’s Group (NIDDK, NIH) and the Hafner Group for valuable comments and suggestions. Specially, Dimitrios Anastasakis (Past Hafner Group member, present University of Crete, Greece) for the source-code of ‘paralus’, Jeremy Scutenaire (Hafner Group) for running the tool and helping identify bugs, and Franziska Ahrend (Haase Group) for reviewing the code. We acknowledge the NIAMS HPC cluster and the NIH HPC Biowulf cluster for computational resources.

## Author contributions

A.H.P. conceived the project, developed the computational method under supervision of M.H.; A.H.P. wrote the manuscript and edited together with M.H.

## Data and code availability

MATR3 PAR-CLIP data (GSE262647) have been deposited in The NCBI Gene Expression Omnibus (https://www.ncbi.nlm.nih.gov/geo/) under the indicated accession numbers. Additional DHX36 and HNRNPK PAR-CLIP data has been downloaded from SRR6191036 (GSE105175) and SRR23199821 (GSE223603) respectively. PCLIPtools is available on GitHub (https://github.com/paulahsan/pcliptools) and Zenodo (https://doi.org/10.5281/zenodo.17064150).

