## Supplementary Fig for "PCLIPtools: A Robust Framework for Identifying RNA-Protein Interaction Sites from PAR-CLIP experiments"

**Authors**
Ahsan H. Polash and Markus Hafner
**Affiliation:**
RNA Molecular Biology Laboratory, National Institute of Arthritis and Musculoskeletal and Skin Diseases (NIAMS), National Institutes of Health (NIH), Bethesda, 20892 Maryland, USA

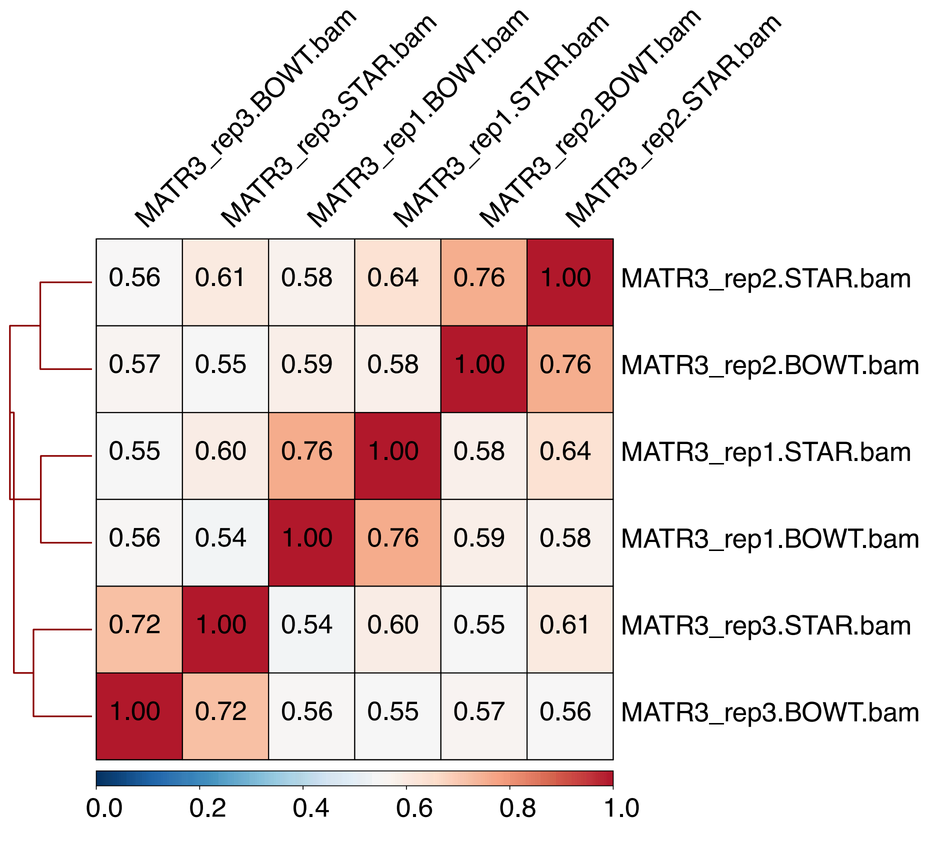


**Supplementary Figure 1.** Both STAR and Bowtie (BOWT) aligners resulted in comparable alignments. However, STAR aligned bam files are more uniform accross replicates.

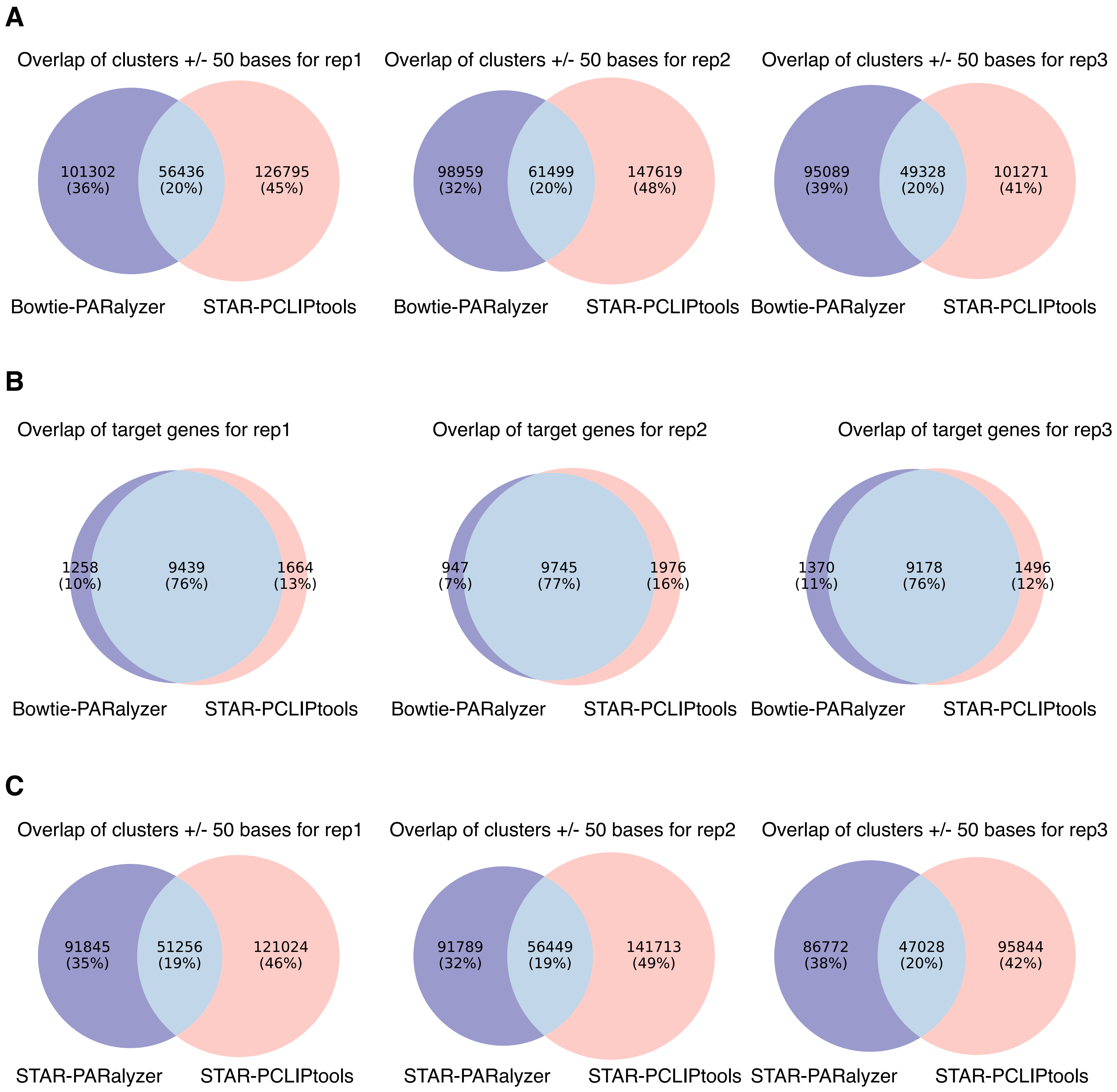


**Supplementary Figure 2.** Comparison of cluster coordinates (+/- 50 bases) for (**A**) Bowtie-PARalyzer and STAR-PCLIPtools, (**C**) STAR-PARalyzer and STAR-PCLIPtool showed substantial amount of clusters were unique due to the differences in the peak calling tools and the aligner tool used. (**B**) Regardless of the alignment method and the peak calling method, comparison of the PAR-CLIP target genes are more uniform than comparing the exact interaction sites.

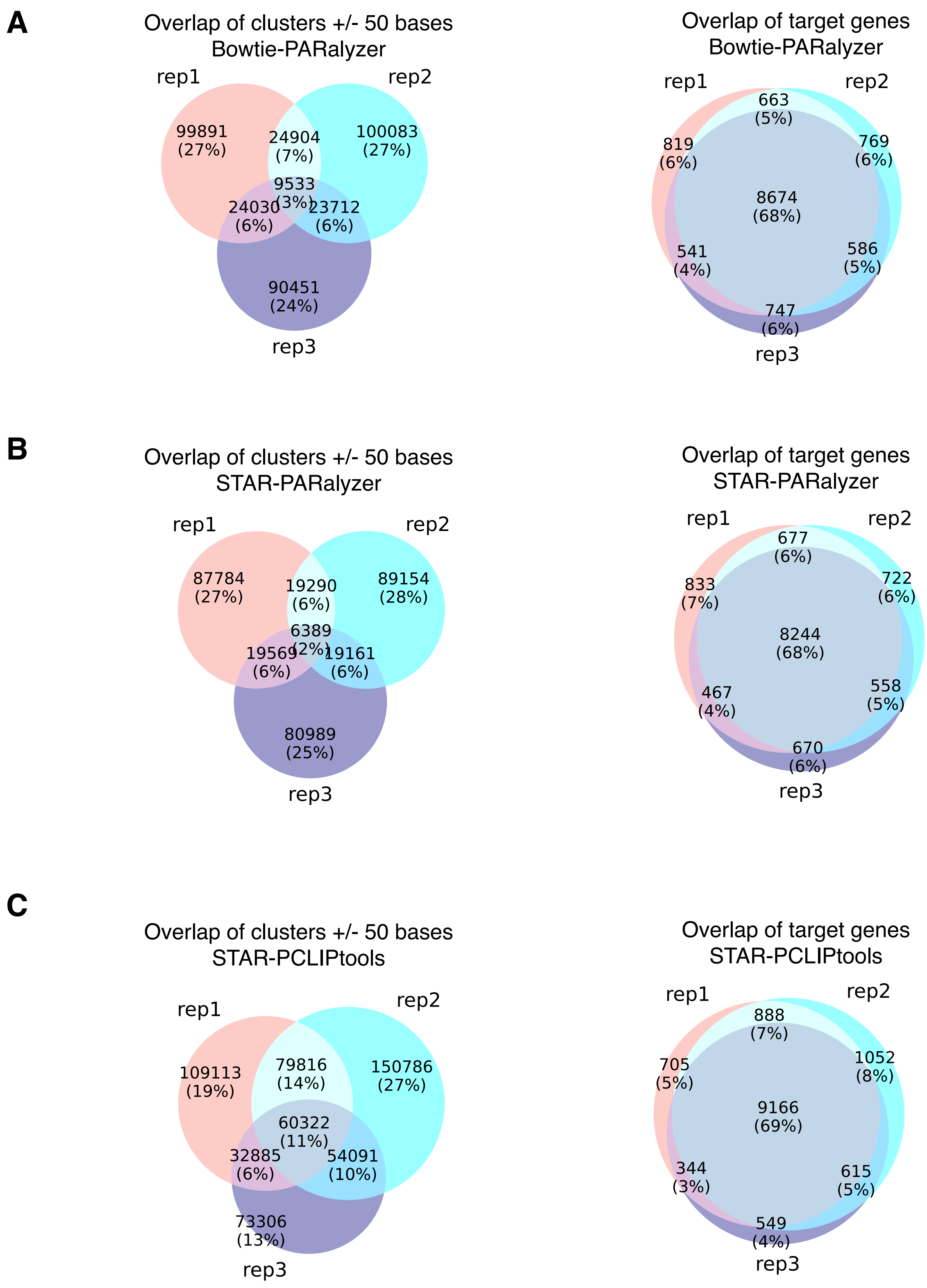


**Supplementary Figure 3.** Comparison of cluster coordinates (+/- 50 bases) derived from (**A**) Bowtie-PARalyzer, (**B**) STAR-PARalyzer and (**C**) STAR-PCLIPtools showed the latter resulted in higher reproducibility accross replicates compared to rest.

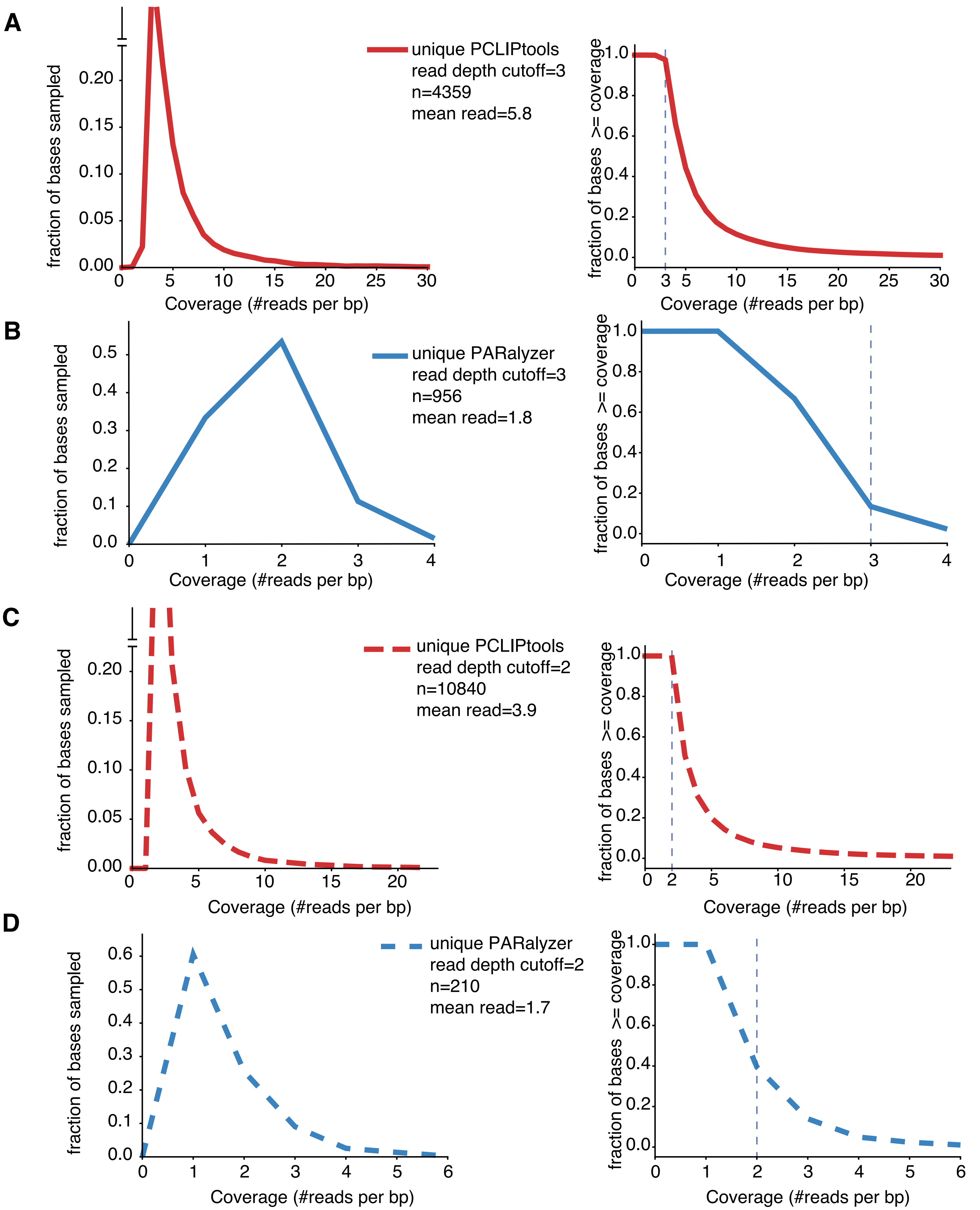


**Supplementary Figure 4:** PCLIPtools specific clusters for chromosome 13 are enriched with high read depth (**A** left panel) and (**C** left panel) comapred to PARalyzer spefic clusters (**B** left panel) and (**D** left panel). PCLIPtools maintains minimum read threshold 3 reads (**A** right panel) and 2 reads (**C** right panel) for clusters whereas PARalyzer often retains clusters that do not meet minimum read depth threshold (**B** right panel) and (**D** right panel).
